# Knockdown of endogenous circulating C1 inhibitor induces neurovascular impairment, neuroinflammation and cognitive decline

**DOI:** 10.1101/216531

**Authors:** Dorit Farfara, Emily Feierman, Allison Richards, Alexey S. Revenko, Robert A. MacLeod, Erin H. Norris, Sidney Strickland

## Abstract

Plasma proteins and activated immune cells are known contributors of vascular brain disorders. However, the mechanisms and routes involved are still unclear. In order to understand the cross-talk between plasma proteins and the brain, we knocked down circulating C1 inhibitor (C1INH) in wild-type (WT) mice using antisense-oligonucleotide (ASO) technique and examined the brain. C1INH is a plasma protein inhibitor of vascular inflammation induced by activation of the kallikrein-kinin system (KKS) and the complement system. This knockdown induced the activation of the KKS but spared the activation of the classical complement system. Activation of the KKS induced an upregulation of the bradykinin pathway in the periphery and the brain, resulting in hypotension. Blood-brain barrier (BBB) permeability, plasma protein extravasations, activated glial cells and elevated levels of IL-1beta, IL-6, TNF-alpha, and iNOS were detected in brains of C1INH ASO treated mice. Infiltrating innate immune cells were evident, entering the brain through the lateral ventricle walls and the neurovascular units. The mice showed normal motor functions, however, cognition was impaired. Altogether, our results highlight the important role of regulated plasma-C1INH as a gatekeeper of the neurovascular system. Thus, manipulation of C1INH in neurovascular disorders might be therapeutically beneficial.

## Introduction

C1 inhibitor (C1INH) is a circulating plasma protein, belonging to the superfamily of serine protease inhibitors (serpins). It circulates in the plasma at the concentration of 0.15–0.3mg/ml and is mainly produced by the liver. It inhibits the activation of the complement system [1] and the kallikrein-kinin system (KKS) [2] which share many inflammatory features mediated by the vascular system [3,4].

Activation of the KKS through plasma kallikrein and high molecular weight kininogen (iHK) induces the secretion of a potent 9-amino acid peptide, bradykinin [5]. Bradykinin is known to cause vasodilation, reduce blood pressure, increase vascular permeability and cell recruitment and induce pro-inflammatory response through activation of the bradykinin receptors [6,7]. In the CNS, bradykinin is a known neuromodulator involved in various inflammatory responses and blood-brain barrier (BBB) permeability [8,9]. The correlation between C1INH and bradykinin was shown in a knockout mouse model of C1INH which induced peripheral vascular permeability through the activation of bradykinin 2 receptor [10]. However, there is no evidence correlating C1INH regulation and its effect on the brain.

The BBB is the first immune gate which maintains a homeostatic environment for resident brain cells such as neurons and glia. The BBB is formed mainly by endothelial cells and tight junction proteins in collaboration with astrocytes, pericytes and microglia/macrophage cells, which separates the blood from the brain. Evidence of plasma proteins in the brain suggests leakage and impairment of the BBB, which contributes to chronic neuroinflammatory and autoimmune disorders such as Alzheimer’s disease, Parkinson disease and multiple sclerosis [11,12].

Neuroinflammation is involved in many immune pathways [13] and is tightly connected to the vascular system [14]. Once induced by an immune trigger, activated glia cells, such as astrocytes and microglia, secrete cytokines, chemokines, and other recruiting signals [15]. These released mediators locally diffuse into the bloodstream, attracting leukocytes to the site of inflammation and upregulating the expression of cellular adhesion molecules, which are necessary for attachment and transmigration across post-capillary venules[16]. It is suggested that recruited of peripheral immune cells can be beneficial or harmful, depending on the immune trigger [17]. Nevertheless, It is agreed that acute inflammation is crucial for protection and repair, as opposed to chronically activated inflammation, which might lead to toxic effects.

Based on this information, we hypothesized that by long-term depletion of endogenous circulated C1INH, we might induce neuroinflammatory effects on the brain via the neurovascular system. To test this hypothesis, we knocked down the expression of circulating C1INH using antisense oligonucleotide (ASO) and examined the brain for neurovascular impairments and neuroinflammation.

## Results

**Knockdown of circulating C1INH activates the kallikrein-kinin system independent from FXII, to produce bradykinin and induce hypotension**. Based on the work of Bhattacharjee et al. [18] demonstrating the efficacy of an antisense oligonucleotide (ASO) knockdown technique targeting circulating C1INH (C1INH ASO), we subcutaneously administered C1INH ASO and scrambled control ASO (CTRL ASO) to ten-weeks-old C57/Bl6J. After twelve weeks of treatment, we determined the levels of circulating C1INH protein expression in the plasma of treated mice and confirmed its depletion (Fig. 1 A and B). Since C1INH inhibits the activation of the kallikrein-kinin system (KKS) through high molecular weight kininogen (iHK), we observed a 50% reduction in levels of iHK (110kDA and 82kDa; iHK-ΔD5, lacking domain 5), in plasma from C1INH ASO compared to CTRL ASO treated mice (Fig. 1A and B), indicating an increase in HK cleavage, as previously suggested [19]. Similar results were obtained in non-reduced conditions (Supp. Fig. 1A-D). Plasma kallikrein cleaves iHK, therefore we examined its levels and found significant increased levels in plasma from C1INH ASO-treated mice compared to controls (Fig. 1A and B). We confirmed this occurrence in a different strain model - C57/C3H, to ensure that these results are not strain-specific (Supp. Fig. 1E-H). To confirm these results seen in western blots indicating iHK cleavage, we measured kallikrein activity using a chromogenic substrate assay [19] and found significantly increased activity of kallikrein by 48% in plasma from C1INH ASO-treated mice compared to plasma from control treated mice (Fig. 1C and D). The traditional activation of the KKS is initiated by activation of the contact system through factor12 (FXII) [19]. However, independent cleavage of iHK from FXII can accrue directly by pre-kallikrein, and C1INH was shown to inhibit the pre-kallikrein cleavage of iHK[20]. Interestingly, in our C1INH ASO treated mice, FXII expression did not differ, which led us to think of an independent mechanism of KKS activity induced by the depletion of C1INH protein. To determine if the activation of the KKS is FXII-dependent, we administered C1INH ASO to FXII knockout mice from the same background using the same concentrations and time duration used for the WT mice. In this knockout model treated with C1INH ASO compared to CTRL ASO, C1INH and iHK protein expressions were significantly reduced whereas plasma kallikrein was increased, similarly to the results obtained with WT mice (Fig. 1E and F). Furthermore, the expression of FXI, a protease cleaved by FXIIa, did not differ in the plasma of WT treated groups, supporting the FXII-independent activation of the KKS.

**Fig. 1.**
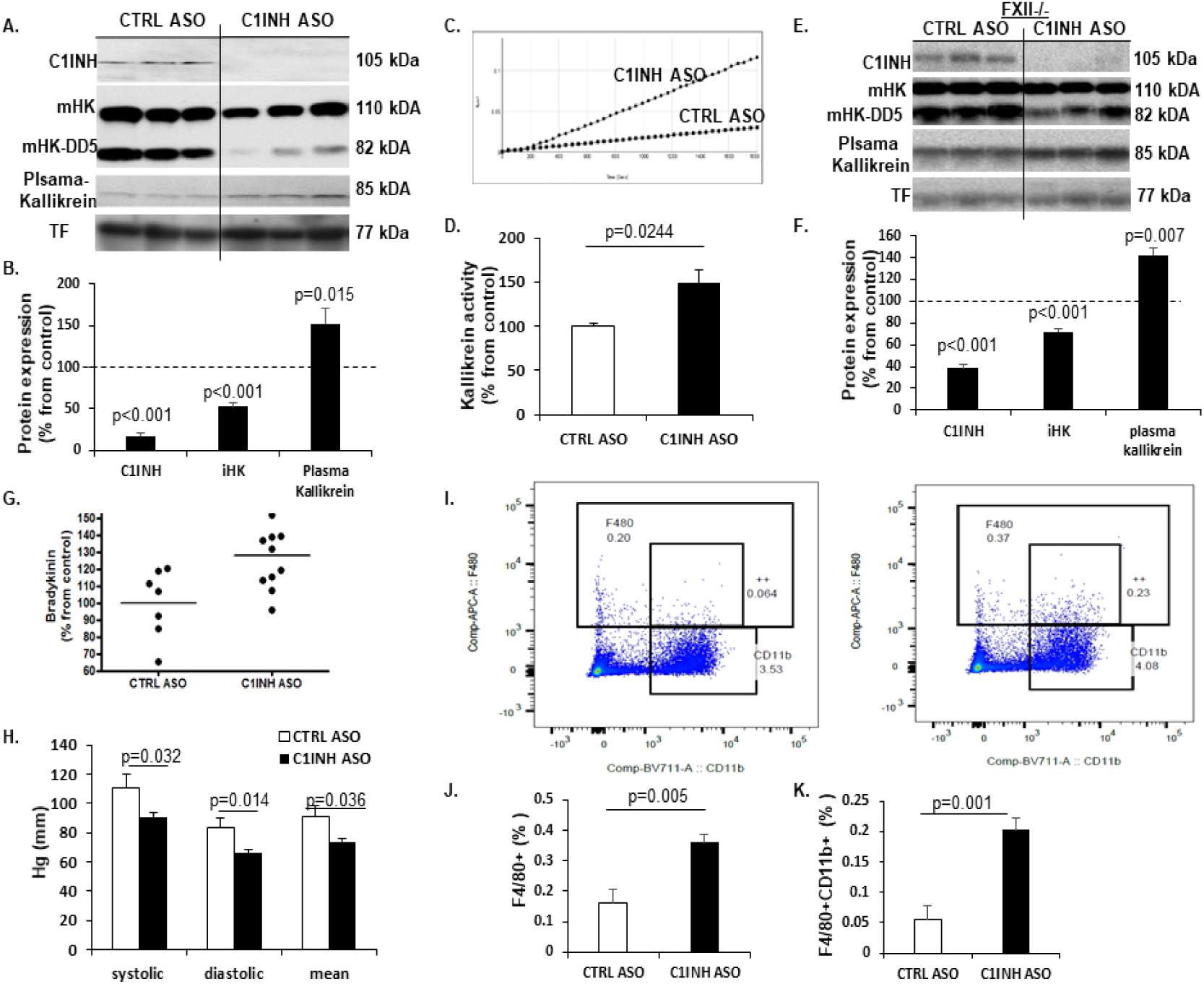
Increased KKS activation, bradykinin production and hypotension in C1INH ASO treated mice. **(A)** Representative Western blots of plasma from mice treated with CTRL ASO vs C1INH ASO, probed with anti-C1INH, ant-intact HK (iHK; 110 kDa and 85 kDa), anti-plasma kallikrein and anti-transferrin antibodies, under reduced conditions. **(B)** Quantification of western blots of plasma from CTRL ASO vs. C1INH ASO showed a significant decrease of 83% in C1INH expression (100±8.79 vs. 17.26±4.27, p<0.001, n=17-19/group) and 55%in iHK expression (100±9.08 vs. 52.72±4.18, p<0.001, n=17-19/group) and increased expression of plasma kallikrein by 52% (100±10.7 vs. 152.3±17.62, p=0.015, n=11-13/group). **(C)** Representative kallikrein activity plot and **(D)** quantification of plasma treated with CTRL ASO vs. C1INH ASO showing a significant increase of 49% in KKS activity in C1INH ASO (100±3.72 vs. 148.73±15.12, p=0.0244, n=5-6/group). **(E)** Representative Western blots of plasma from FXII-/- mice treated with CTRL ASO vs C1INH ASO, probed with anti-C1INH, anti-intact HK (iHK; 110 kDa and 85 kDa), anti-plasma kallikrein and anti-transferrin antibodies, under reduced conditions. **(F)** Quantification of western blots of plasma from CTRL ASO vs. C1INH ASO showed a significant decrease of 62% in C1INH expression in plasma of FXII-/- (100±8.8 vs. 38.29±3.5, p<0.001, n=4-5/group) and 29% reduction in iHK (100±2.32 vs. 71.66±3.35, p<0.001, n=4-5/group) and increased expression of plasma kallikrein by 42% (100±6.62 vs. 141.97±6.64, p=0.007, n=4-5/group) **(G)** ELISA analysis showing 28% significantly more bradykinin in the plasma of C1INH ASO treated mice compared to CTRL ASO group (128±5.85 vs. 100±7.6, p=0.009, n=7-11/group). **(H)** Blood pressure measurement of CTRL ASO vs. C1INH ASO of systolic (111.18±9.05 vs. 90.8±2.97, p=0.032) diastolic (83.49±6.49 vs. 65.81±2.63, p=0.014) and mean (90.85±7.74 vs. 73.83±2.59, p=0.036) indicate that C1INH ASO treated mice are hypotensive vs. CTRL ASO (n=9-11/group). **(I)** Representative FACS dot-plot of positive F4/80 vs. CD11b from peritoneal macrophages of CTRL ASO vs. C1INH ASO treated groups. **(J)** Quantification analysis of F4/80 positive cells significantly increased in peritoneal macrophages from C1INH ASO compared to CTRL ASO (0.161±0.043 vs. 0.36±0.02, p=0.005, n=5/group). **(K)** Quantification analysis of F4/80 and CD11b positive cells (F4/80+CD11b+) significantly increased in peritoneal macrophages from C1INH ASO compared to CTRL ASO (0.054+0.02 vs. 0.2+0.019, p=0.001, n=5/group).

The cleavage of iHK yields a 9-amino acid peptide called bradykinin. Bradykinin is unstable in the plasma and is rapidly eliminated by degradation. Using a sensitive ELISA kit for detecting bradykinin, we found that levels of bradykinin were significantly increased in C1INH ASO-treated mice compared to CTRL ASO-treated mice (Fig. 1G). LPS injected mice were used as positive controls (Supp. Fig. 1I). Elevated levels of bradykinin are known to correlate with low blood pressure [21,22,23] and indeed, we found a significant decrease in systolic, diastolic, and mean blood pressure in C1INH ASO-compared to CTRL ASO-treated mice (Fig. 1H).

We next confirmed the activation of peritoneal macrophages by bradykinin activation pathway, as previously suggested [24]. By measuring levels of F4/80 and CD11b via FACS, we determined a significant increased activation of peritoneal macrophages in C1INH ASO-treated mice compared to CTRL mice (Fig. 1I-K), suggesting that the depletion of C1INH had induced activation of peritoneal macrophages. We also checked levels of C1INH in this macrophages population and found it to be significantly increased by 64% (Supp. Fig. 2A and B), suggesting the ASO treatment targeted only the circulated protein and did not affect the protein expression in macrophages.

**Fig. 2.**
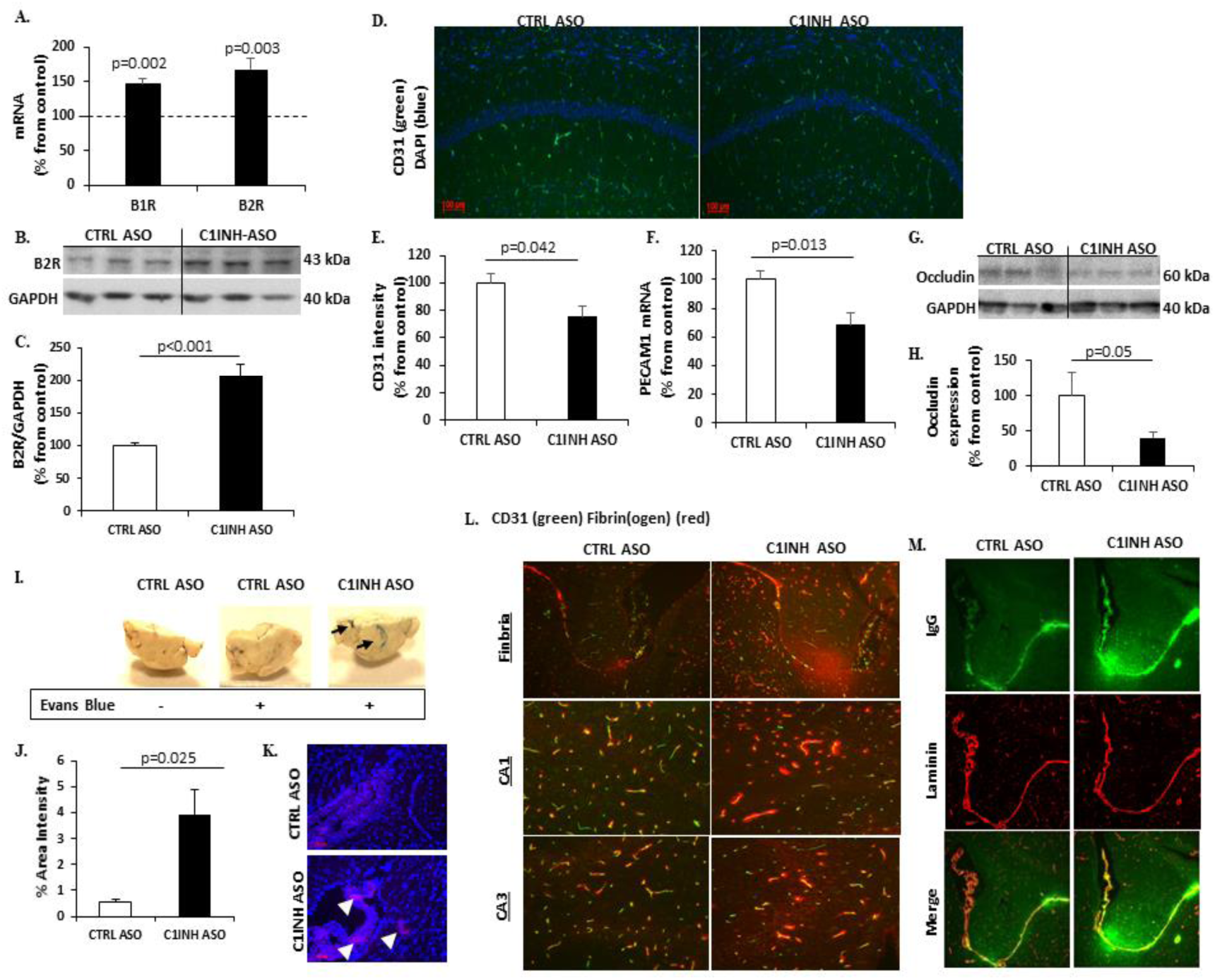
Knockdown of circulated C1INH leads to activation of the bradykinin pathway and BBB permeability. **(A)**. RTPCR of coronal brain samples showed elevated gene expression of B1R (100±6.86 vs. 145.99±9.02, p=0.002, n=6) and B2R (100±3.98 vs. 167.44±16.67, p=0.003, n=7-9) from CTRL ASO treated mice compared to C1INH ASO. **(B)** Representative western blot of B2R of coronal brain homogenates compared between treated groups. **(C)** Quantification of B2R expression levels show a 2-fold increase in brain from C1INH ASO compared to CTRL ASO (100±3.95 vs. 206.75±17.85, p<0.001, n=5). GAPDH was used for normalization. **(D)** Representative image of coronal section stained with anti-CD31 and DAPI (bar is 100μm scale). **(E)** Quantification of the intensity of CD31 show a decreased percentage in CA1 region of CTRL ASO and C1INH ASO (100±7.29 vs. 75.67±7.75, p=0.042, n=7-9/group). **(F)** PECAM1 gene expression is decreased in brains of C1INH ASO vs. CTRL ASO (100+6.54 vs. 68.56+8.27, p=0.013, n=6). **(G)** Representative western blot of Occludin expression. **(H)** Quantification of occluding expression showed a decreased signal in C1INH ASO compared to CTRL ASO (100+32.69 vs. 40.64+7.73, p=0.055, n=7-10/group). **(I)** Representative images of brain hemispheres from mice after Evans blue injection and perfusion. **(J)** Quantification of blue area from brain sectioned area in percentage using ImageJ (p=0.025, n=3). **(K)** Representative staining of Evans blue (red) in the brain parenchyma and the lateral ventricle. Blue signal is DAPI staining (bars are 100μm scale). **(L)** Representative image of CA1 and CA3 brain regions stained with CD31 (green) and fibrin(ogen) (red). **(M)** Representative image of IgG (green) and laminin (red) in the lateral ventricle and fimbria between CTRL ASO and C1INH ASO brains.

In order to rule out liver toxicity as a possible outcome of the ASO treatment we examined levels of alanine aminotransferase (ALT) and found no differences between plasma from the treated groups (supplement Fig. 3A). In addition, acquired immune response demonstrated by T cells and B cells activation, and spleen enlargement was also not detected (Supp. Fig. 3B-E). C1INH expression in CD4+ and CD8+ cells, showed no differences between the treated groups (Supp. Fig. 2C-F). We also ruled out stress as an outcome of the chronic ASO treatment and a possible mechanism of immune activation, by measuring corticosterone (CORT) levels in the plasma and saw no differences between treated groups (Supp. Fig. 3F).

C1INH is majorly known as an inhibitor of the complement system, however we found no changes in classical complement activation proteins measured by western blots and ELISA for C1qA, C1r and C3a (Supp. Fig. 4A-G) suggesting that the depletion of circulating C1INH leads to the activation of the KKS without involvement of the activation of the complement system.

**Knockdown of circulating C1INH increased bradykinin receptors in the brain and mediated BBB permeability**. As a potent vasodilator, bradykinin was shown to induce BBB permeability [25,26,27] and mediate pro-inflammatory response in the nervous system through its receptors [28]. When examining the gene expression of bradykinin 1 receptor (B1R) and bradykinin 2 receptor (B2R) in the brains of C1INH ASO- vs. CTRL ASO-treated mice we saw significant increase by 45% and 67%, respectively (Fig. 2A). Moreover, B2R protein expression was increased by two-folds in cortex of C1INH ASO mice compared to controls (Fig. 2B and C), suggesting an elevated activation of the bradykinin pathway in the brain.

Next, we examined endothelial cells expression as the major components of the BBB. Using CD31 staining we showed a 25% decreased expression in the hippocampus of the CA1 region (Fig. 2D and E) in C1INH ASO- vs. CTRL ASO-treated mice. Gene expression of PECAM1 from C1INH ASO brains showed a 32% down-regulation levels compared to CTRL ASO brains (Fig. 2F). Occludin, a tight-junction transmembrane protein of brain endothelial cells which has been shown to be degraded with increased BBB permeability[29], was also decreased by 60% between brains from C1INH ASO and CTRL ASO-treated mice (Fig. 2G and H).

At the end of the chronic treatment we injected Evans-Blue to CTRL ASO- and C1INH ASO-treated mice to determine BBB permeability. We found an increase in blue staining in the freshly frozen brains of C1INH ASO mice, specifically in the ventricles and the interstitial spaces (Fig. 2I and J). Moreover, Evans blue dye was detected by fluorescence microscopy in the brain tissue of C1INH ASO mice but not CTRL ASO, specifically in the margins of the lateral ventricle (Fig. 2K). In addition, fibrin(ogen) and IgG, which are peripheral components, detected in the brains only in barrier-impairment conditions, showed extensive extravasation from blood vessels in the region of the lateral ventricle and the fimbria, CA1, and CA3 regions in brains of C1INH ASO compared to CTRL ASO-treated mice (Fig. 2L and M), supporting the findings of BBB disruption. When we analyzed the whole brains for differences in water content, to test for brain edema, we did not find any changes between treated groups (Supp. Fig. 5).

**Increased activation of resident glia cells towards a pro-inflammatory response**. Astrocytes and microglia cells are the resident immune cells of the brain and along with neurons and vascular cells they comprise the neurovascular unit (NVU) [30]. Upon activation they secrete pro- and anti-inflammatory cytokines depending on the immune trigger [15], which contribute to BBB permeability and infiltration of peripheral immune cells [31]. We examined astrocyte activation by analyzing GFAP levels and activated macrophage/microglia using CD11b levels, which are known markers known to be upregulated in activation. We found significant increased up-regulation of both genes (Fig. 3A) and proteins expression of GFAP and CD11b (Fig. 3B and C) in the cortex of C1INH ASO treated mice compared to CTRL ASO mice. We also confirmed the results using immunofluorescence staining for astrocytes and microglia/macrophages (Fig. 4D).

**Fig. 3.**
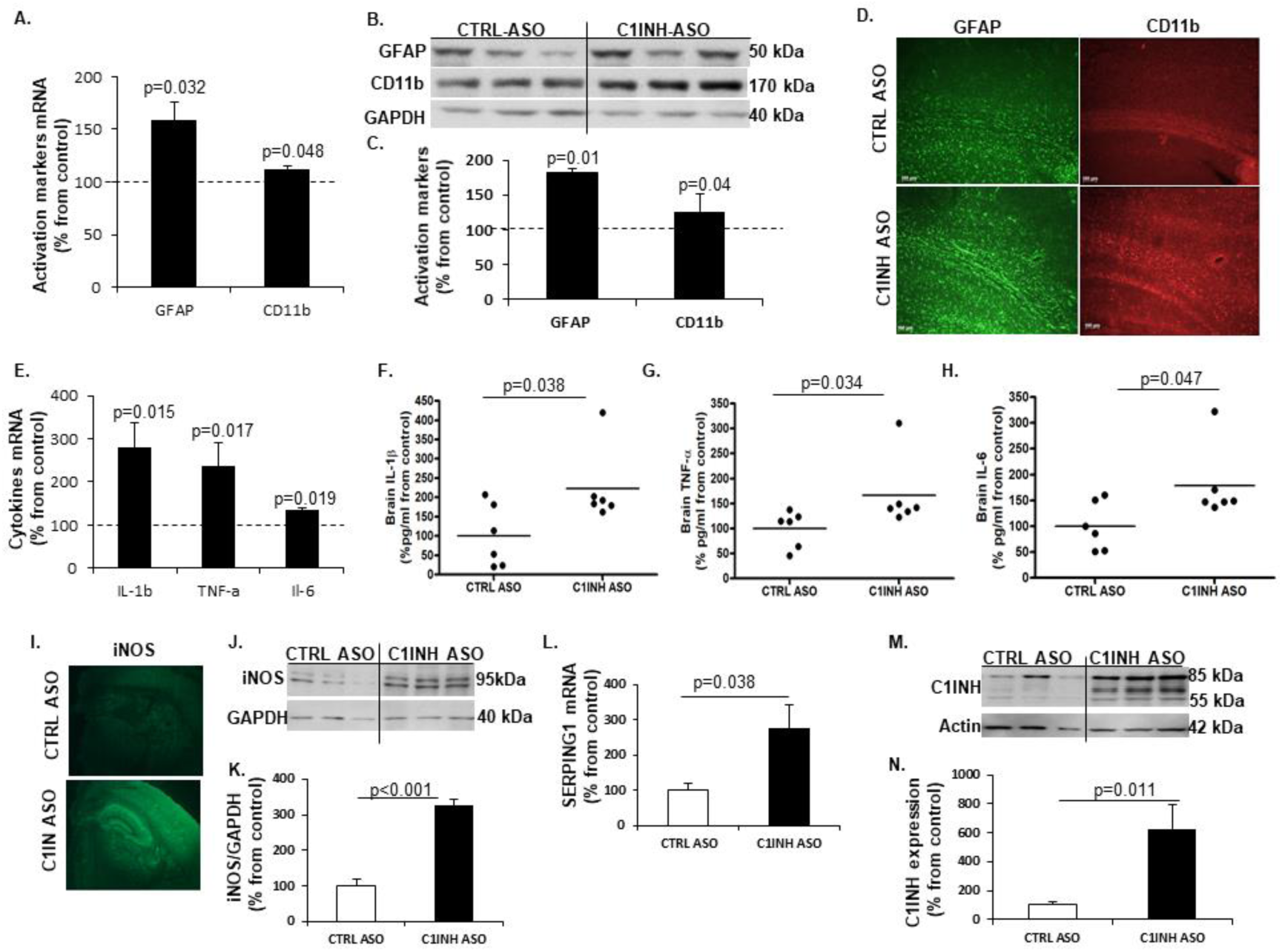
Glial cell activation towards a pro-inflammatory response. **(A)** Increased gene expression of GFAP (158.57+17.2 vs. 100+11.69, p=0.032; n=4-10/group) and CD11b (112.54+3.14 vs. 100+7.36, p=0.048, n=4-10/group) in the cortex of mice treated with C1INH ASO compared to control treatment. **(B)** Representative Western blot of GFAP and CD11b in the cortex of C1INH ASO treated mice vs. CTRL ASO treated mice. **(C)** Increased protein expression of GFAP (181.22+25.19 vs. 100+7.14, p=0.014, n=5) and CD11b (126.3+10 vs. 100+3.94, p=0.04, n=5) in cortex of C1INH ASO treated mice compared to CTRL ASO treated mice. **(D)** Representative immunofluorescence staining of GFAP and CD11b in the cortex of C1INH ASO vs. CTRL ASO treated mice (100μm scale). **(E)** Elevated gene expression of IL-1β (279.34+58.16 vs. 100+20.52, p=0.015, n=5-7/group), TNF-α (236.15+54.32 vs. 100+20.25, p=0.017, n=5-7/group) and IL-6 (133.70+7.06 vs. 100+12.9, p=0.019, n=5-7/group) in whole brains of C1INH ASO treated mice compared to CTRL ASO treated mice. Increased protein expression levels of **(F)** Il-1β (223.2+39.66 vs. 100+33.02, p=0.038, n=6), **(G)** TNF-α (166.4+29.09 vs. 100+14.87, p1=0.034, n=6) and **(H)** IL-6 (178.8+29.09 vs. 100+19.18, p=0.047, n=6) in C1INH ASO treated mice vs. control group. **(I)** Representative immunofluorescence staining of iNOS in the brains of C1INH ASO treated mice compared to CTRL ASO treated mice. **(J)** Representative Western blot of iNOS in brain homogenates of C1INH ASO treated mice compared to controls. **(K)** Elevated protein expression of iNOS (326.32+18.64 vs. 100.1+18.52, p<0.001, n=6) in C1INH ASO treated mice compared to controls. **(L)** Increased levels of SEPING1 gene expression in whole brains of C1INH ASO compared to CTRL ASO (227.4+46.78 vs. 100+13.36, p=0.038, n=6). **(M)** Representative western blot of C1INH of whole brain homogenates compared between treated groups. **(N)** Increased protein expression of C1INH in whole brains of C1INH ASO vs. CTRL ASO (62732+170.46 vs. 100+23.86, p=0.011, n=6).

Activation of glial cells is known to mediate pro-inflammatory cytokine production [15]. We first measured mRNA profile of pro-inflammatory cytokines such as IL-1β, TNF-α and IL-6, since they are the major secreted cytokines from activated glial cells in chronic neuroinflammatory disorders. Comparing brain samples from C1INH ASO- and CTRL ASO-treated mice, RT-PCR analysis confirmed the elevation in pro-inflammatory cytokine gene expression (Fig. 3E) and ELISA confirmed an increase in their proteins levels (Fig. 3F-H). Furthermore, since nitric oxide is known to be secreted from activated resident immune cells during pro-inflammatory response [15], we used immunofluorescence staining (Fig. 3I) and western blotting to examine changes in inducible nitric oxide synthase (iNOS). We observed a significant increase in iNOS expression in brains of C1INH ASO mice compared to CTRL ASO-treated mice (Fig. 3J and K). Interestingly, C1INH levels in the brain were highly elevated both in gene levels and protein expression (Fig. 3L-N) indicating that the ASO treatment did not affect levels of C1INH in the brain and that the elevation seen is might be a result of a compensatory mechanism towards increased pro-inflammatory response. To ensure the lack of involvement of the classical complement system as seen in the periphery, we examined the levels of C3a and C3aR1 in the brains of treated mice using RTPCR and ELISA, and found no differences (Supp. Fig. 6A and B).

**Knockdown of circulating C1INH increased levels of infiltrating myeloid cells in the brains of C1INH ASO-treated mice**. The brain is constantly monitored by resident and non-resident innate immune cells. The circumventricular organs (CVO’s) are located around the ventricles and are known areas of interaction between the blood, the CSF, and the brain parenchyma. Due to their unique structure, the passage of large substances and cells from the blood to the perivascular spaces and parenchyma is evident, exposing the brain to peripheral signaling[32]. The Choroid plexus is thought to be part of the CVO’s because of its fenestrated capillaries. The CP in the ventricular system and the meningeal vasculature is agreed to be a selective gate for transmigrating immune cells into the brain [33]. As a multifunctional organ, the choroid plexus (CP) which produces the CSF, is abundant with myeloid cells that function as immunosurveillance cells in health and disease [34]. Infiltrated myeloid cells exhibit enhanced phagocytic capacity, neurotrophic support and anti-inflammatory characteristics compared to resident microglia [35,36]. Resident microglia cells are distinct from peripheral macrophages that infiltrate the CNS under pathological conditions by origin and function [37]. Recent findings have distinguished between resident microglia from infiltrating macrophages using TMEM119 [38] and significant morphology differences [35]. In order to evaluate the immune response in the brain as a result of C1INH knockdown, we used CD11b, CD68 which is expressed highly by phagocytic macrophages and activated microglia cells, along with TMEM119 staining. We identified resident microglia cells as those co-localizing with CD11b and TMEM119 (CD11b+/TMEM119+) vs. non-resident monocyte-derived cells which express CD68 or CD11b but not TMEM119 (CD68+/TMEM- or CD11b+/TMEM-). We examined the general population of cells in the lateral ventricle (LV) and among the CP cells of WT CTRL mice, and found intense staining of CD68 and CD11b cells, which are negative for TMEM119, confirming the published data indicating blood-derived macrophages cells populate the ventricles. Moreover, the morphology of CD68+/TMEM- or CD11b+/TMEM- cells observed in the lateral ventricle, were defined by large cell bodies with small process as opposed to resident microglia cells expressing CD11b+/TMEM119+ surrounding the lateral ventricle which are detected by smaller cell bodies and longer processes (Fig. 4A). Along with GFAP staining for astrocytes detection, we determined that most of the high expressed CD68 cells were either in the lateral ventricle or bordered by GFAP around cerebral blood vessels (Supp. Fig. 7), supporting the evidence that those cells are not resident immune cells. PVM are mostly located in the perivascular spaces surrounding arteries and veins throughout the brain tissue[39] and are elevated in brain disorders[39]. In order to distinguish between blood-derived infiltrating macrophages and perivascular macrophages (PVM) we used CD206 co-stained with CD68 (CD206+/CD68+). We identified positive CD68 which co-localize with CD206 in the PV spaces and the LV of WT CTRL brains, all negative to TMEM119, suggesting these cells are not residential but perivascular macrophages (Supp. Fig. 8).

**Fig. 4.**
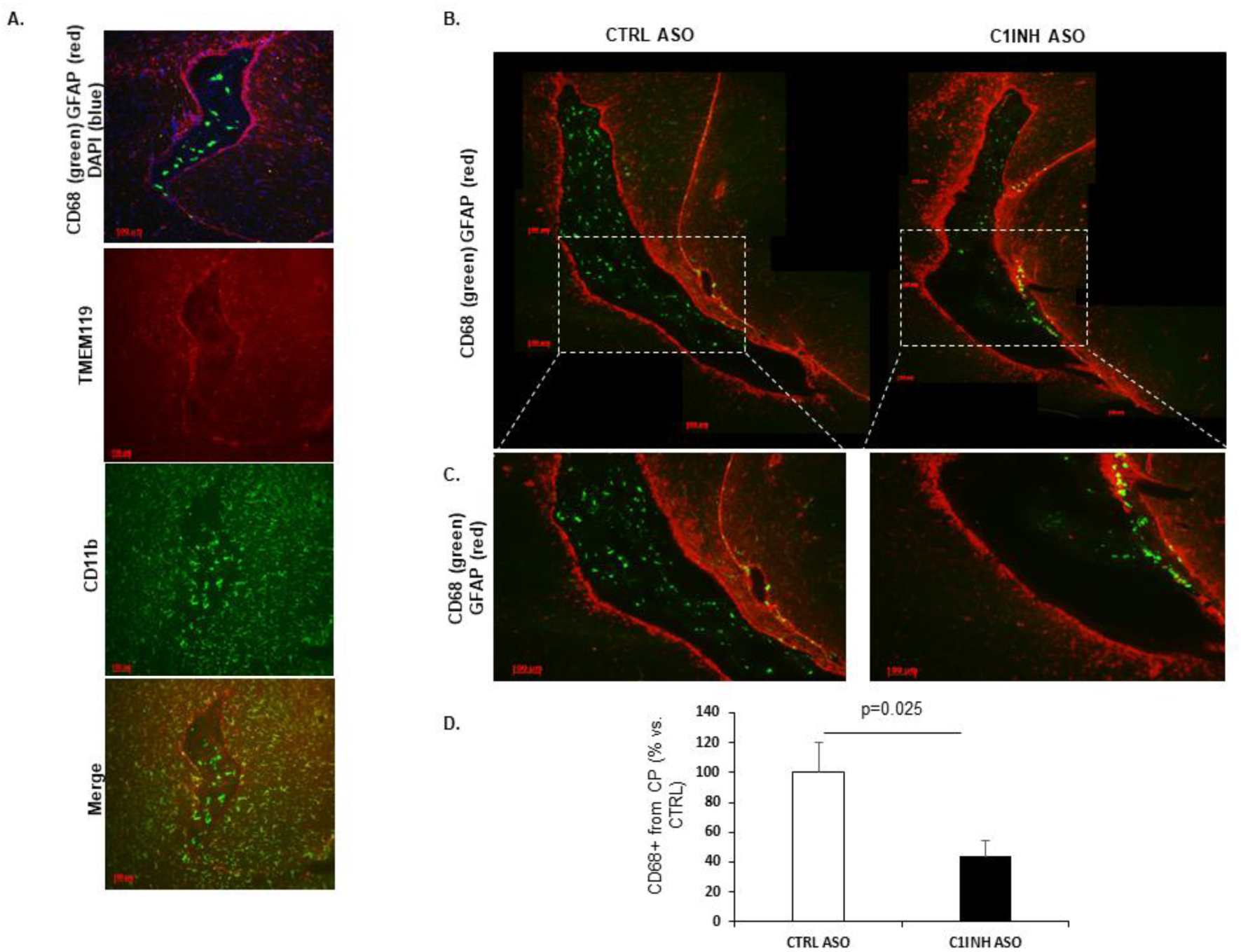
Infiltrating blood-derived macrophages through ventricular spaces. **(A)** Representative images of the lateral ventricle of CTRL ASO mice stained with CD68 (green), GFAP (red) and DAPI (blue). CD11b cells were positively stained in the lateral ventricle and negatively stained for TMEM119. **(B)** Representative stitched images of the lateral ventricle in bregma −2.3mm (approximately) stained with CD68 (green) and GFAP (red) between CTRL ASO and C1INH ASO treated mice. **(C)** Representative images zooming x10 in the lateral ventricle shown in B, showing migrating CD68 positive cells in the lateral ventricle wall area in C1INH ASO treated mice brains. **(D)** Quantification of CD68 by intensity of staining in CTRL ASO vs. C1INH ASO treated mice brains (100±20.19 vs. 43.94±10.45, p=0.025, n=9). (bar scales are 100μm scale)

Based on this characterization, Fig. 4B-D shows a 56% reduced expression of CD68 positive cells with round-shaped bodies, identified as non-residential macrophages, surrounding the CP and in the lateral ventricle of C1INH ASO treated mice compared to CTRL ASO mice. In contrary, high expression of CD68 positive cells, were evident in the walls of the lateral ventricles and the perivascular spaces of C1INH ASO treated mice compared to CTRL ASO (Fig. 4D), suggesting recruitment of non-residential macrophages to the brain through the vasculature.

Moreover, in the brains of C1INH ASO treated mice we saw more cells that were positive to CD68 cells and negative for CD206 staining in the lateral ventricle wall and in different areas of the parenchyma and white matter such as the optic tract (Fig. 5), suggesting increased infiltration of non-residential macrophages which are not perivascular origin in the brains of C1INH ASO treated mice.

**Fig. 5.**
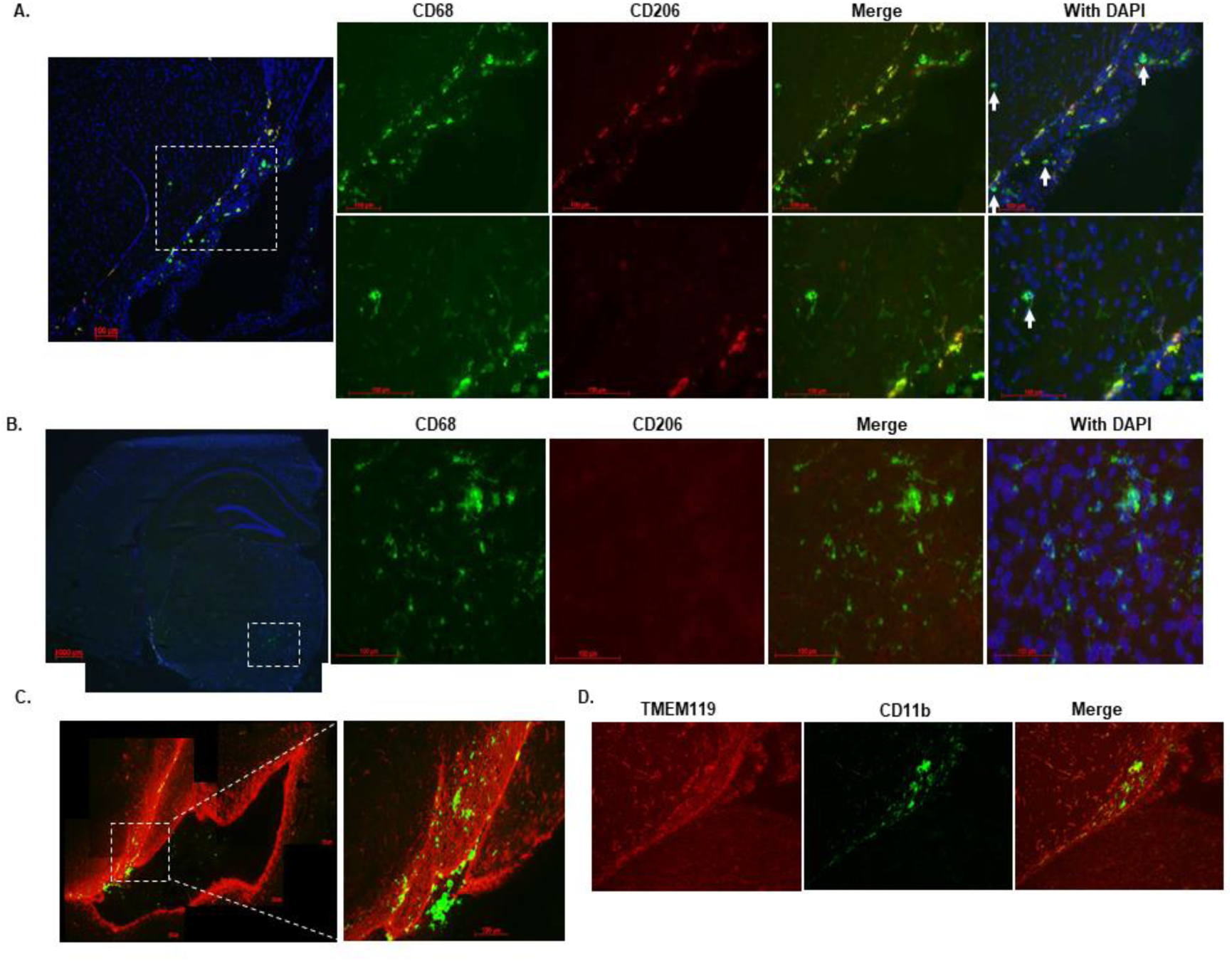
Infiltrating blood-derived macrophages in different regions of the brain. **(A)** Representative image of the ventricular wall stained with CD68 (green), CD206 (red) and DAPI (blue) showing infiltrating blood derived cells that are not co-localized with CD206 (white arrows) **(B)** Representative image of positive CD68 cells that are not co-localizing with CD206 in the brain parenchyma. **(C)** Representative image from bregma −2.54mm of C1INH ASO treated mice brain showing migrating CD68 positive cells in the optic tract region. **(D)** TMEM119 (red) was negatively stained whereas CD11b (green) was positively stained in the optic tract of C1INH ASO treated mice brain (bar scales are 100μm scale).

**Knockdown of circulating C1INH in WT mice result in cognitive decline**. At the end of the chronic treatment, we performed behavioral tests to evaluate motor and cognitive function in mice. Using the fear conditioning paradigm, C1INH ASO-treated mice showed no difference in percentage of movement in the first day when receiving the electric shock compared to the CTRL ASO group (Fig. 6A), however, they showed a reduced percentage of total freezing time in the second day of the test without the electric shock (Fig. 6B), indicating impairment in learning and cognition. In order to evaluate motor disabilities that might be affected by the ASO treatment, we quantified the average distance traveled on the first day of the fear conditioning test prior to foot shock. No motor dysfunction was observed between the groups (Fig. 6C). We also performed an open field test to compare the total distance the mice traveled after treatment and found no differences between the treated groups (Fig. 6D). Since there was no difference observed in motor function between the C1INH ASO and CTRL ASO groups, the reduced freezing time during fear conditioning is most likely a result of cognitive impairment due to the chronic depletion of peripheral C1INH. Although cognitive impairment is often correlated with neurodegeneration, we did not observe apoptotic cell degeneration measured by TUNEL (Supp. Fig. 9).

**Fig. 6.**
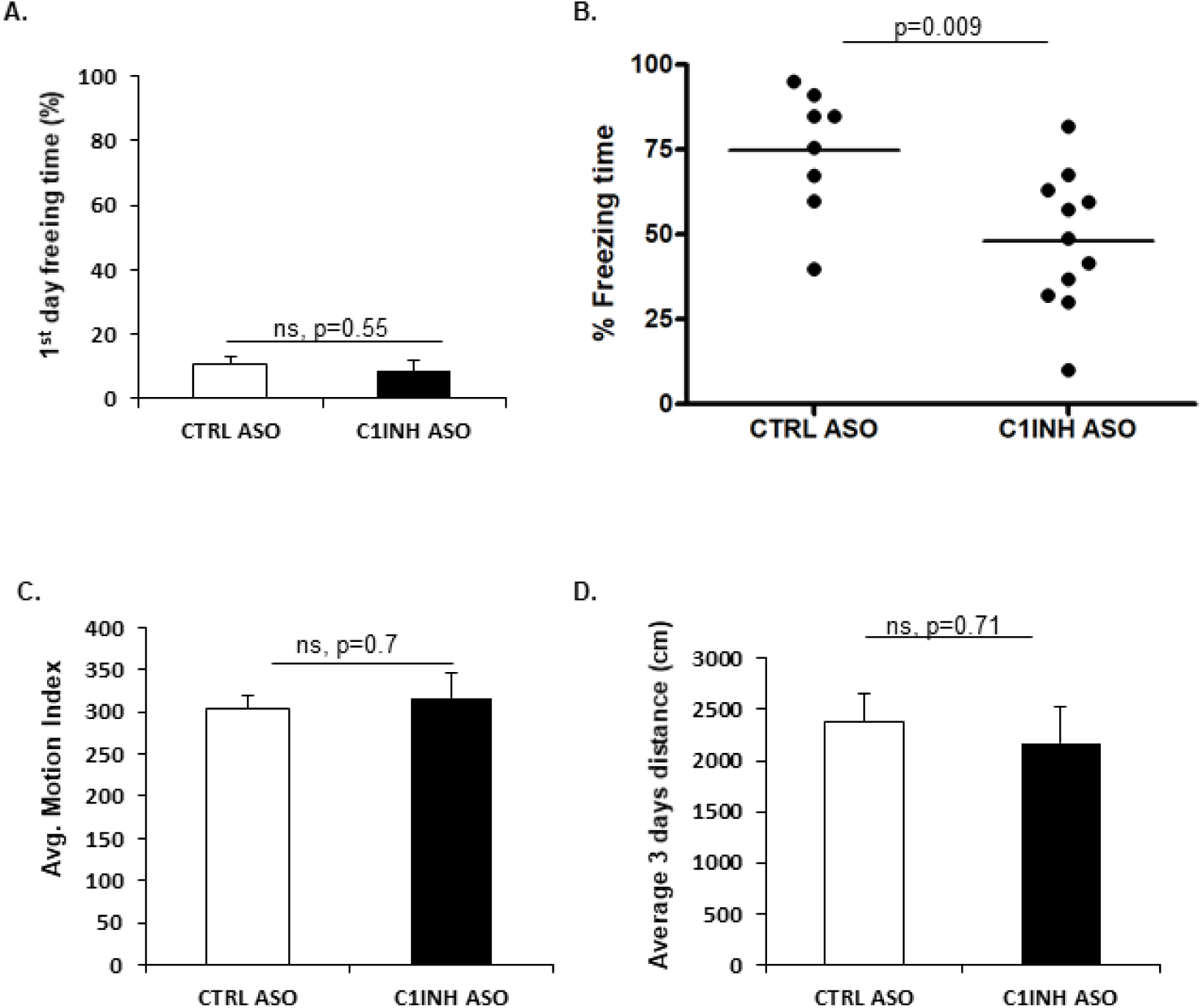
Cognitive dysfunction in C1INH ASO treated mice. **(A**) No differences in the percentage of freezing time on the 1^st^ day of the fear conditioning test between treated groups (10.85+2.02 vs. 8.49+3.3, p=0.55, n=5). **(B)** Decreased freezing time on the 2^nd^ day of the fear conditioning test in C1INH ASO treated mice compared to controls (48.02+6.137 vs. 74.8+6.52, p=0.009, n=8-11/group). **(C)** Average motion analysis of the first day of the fear conditioning test showing no differences between treated groups (304.58+14.22 vs. 317.29+28.7, p=0.7, n=5). **(D)** No differences in measurement of the average of three days distance movement in an open field test (2375.16+180 vs. 2175.53+200, p=0.7, n=3-6/group).

## Discussion

To our knowledge, this is the first demonstration showing that by depleting the endogenous levels of a circulating plasma protein in WT mice, brain inflammation is induced through the vascular system. In this study, we show that chronic depletion of circulating endogenous C1INH by antisense oligonucleotide (ASO) technique, lead to neuroinflammation and cognitive impairment mediated by the vascular system. By depleting the normal levels of C1INH in the plasma of WT mice, we induced the activation of the kallikrein-kinin system (KKS) to produce high levels of bradykinin and induce hypotension. Interestingly, the classical complement system was not activated. After ruling out possible toxic effects in the periphery and an in-direct depletion effect of C1INH expression in different cells and the brain due to the ASO treatment, we analyzed the brain. In the brain, bradykinin pathway was up-regulated, the BBB was permeable, peripheral monocyte-derived cells infiltrated through the vascular system into the brain, pro-inflammatory cytokines were secreted, and cognitive dysfunction was observed.

Our knockdown mouse model partially mimics the clinical pathology of Hereditary angioedema (HAE) type-1, a condition caused by mutations in the SERPING1 gene leading to depletion of C1INH production in humans. HAE type-1 and acquired angioedema (AAE) type-1 are diseases characterized by recurrent episodes of severe, localized inflammation and increased vascular permeability affecting soft tissues, including the gastrointestinal tract, upper airways, and the mucosa [40,41]. Recent evidence has indicated occurrence of migraines, stroke, cerebral symptoms, blindness, tetraspastisity and other irreversible brain damages in HAE patients [42,43,44,45,46]. Similar to our results, plasma from C1INH-deficient patients contains decreased levels of iHK [47]. During inflammatory attacks, HK is cleaved [48] and bradykinin is released [49]. In difference to our results, the inflammatory acute attacks in HAE patients are mediated by the activation of the KKS along with the complement system [3,50]. However, in our knockdown mouse model, the complement system was quiescent, suggesting that the severe acute attacks seen in HAE patients are not induced just by reduced or impaired C1INH protein levels, but by physiological stress known to trigger the attacks [51] that involves the contribution of the complement system and the KKS together. Today, HAE patients are treated with either recombinant C1INH or kallikrein inhibitors [52,53].

In our study, depletion of circulating C1INH led to decreased blood pressure likely due to the activation of the bradykinin pathway in the plasma and brain, which causes blood vessels to dilate. It was shown that hypoperfusion which causes BBB breakdown, induces memory deficits and glial cells activation [54]. We report here that the impairment of the vasculature is evident also in the brains of the C1INH ASO treated mice. We concluded a blood-brain barrier permeability and plasma proteins extravasations in the brain of C1INH ASO-treated mice. The breakdown of the BBB induces activation of the resident glial cells.

Resident microglia and astrocyte cells are the immune cells of the brain, constantly monitoring the brain for harmful changes. Upon activation, they upregulate activation markers and secrete cytokines depending on the immune trigger [15]. We report here that the microglia and astrocyte cells of C1INH ASO-treated mice showed increased activation towards a pro-inflammatory response observed by elevated levels of IL-1beta, IL-6, TNF-alpha, and iNOS in the brains of C1INH ASO-treated mice.

Supporting evidence report that neuroinflammation and cognitive decline correlate with increased peripheral immune cells activation [55,56,57]. Infiltration of peripheral immune cells to the CNS under inflammatory conditions, is mediated by migration through the ventricular wall into the perivascular spaces and across the glia lamitans penetrating the parenchyma[58]. In the brains of C1INH ASO-treated mice we observed accumulated infiltration of non-residential cells to the brain. In the choroid plexus of the lateral ventricle we saw decreased levels of infiltrating non-resident brain macrophages, expressed by positive CD68 cells which were negative to TMEM119. However, these peripheral innate immune cells seemed to infiltrate from the CP to the ventricle wall through the vascular system, to different parts of the brain. These cells were also negative to CD206, a marker expressed on perivascular macrophages, suggesting they are of different origin. We determined a significant activation of peritoneal macrophages in C1INH ASO treated-mice compared to the controlled groups.

It was suggested that blood proteins and neurovascular units are mediators of cognitive impairment [59,60,61]. When we compared cognition between the treated groups, we confirmed a significant impaired cognitive function.

Treatment with C1INH is proven beneficial in a variety of inflammatory diseases such as hereditary angioedema (HAE), sepsis, myocardial ischemia-reperfusion injury and brain injuries [62]. The therapeutic roles of C1INH have been observed in several animal models of peripheral inflammatory disease including sepsis[63,64,65,66], xenograft transplant rejection[67,68], myocardium ischemia-reperfusion injury[69,70], and hemorrhagic shock [71]. Beyond the common clinical treatment with recombinant C1INH for treating HAE, C1INH was shown to be therapeutically beneficial in humans in coronary and myocardial infarction[72,73], septic shock and vascular leak syndrome [74], and other acute inflammatory diseases[75,76]. Moreover, mice infected with *Streptococcus pneumonia* and treated with C1INH showed an improvement in clinical signs of bacterial clearance in the cerebrospinal fluid (CSF) and blood, decreased leukocyte infiltration to the CSF, meningitis, and reduced Il-6 levels [77]. Furthermore, C1INH was shown to interfere with endothelial-leukocyte adhesion [78,79]. Administration of C1INH was shown to have neuroprotective roles in neurodegenerative diseases such as stroke and traumatic brain injury [80,81,82] by reducing the infarct volume and the neuronal damage in the temporal cortex, striatum, hippocampus, and thalamus.

Our data suggest that chronic depletion of circulating endogenous C1INH can cause neurovascular dysfunction, neuroinflammation and cognitive decline mediated by the activation of the KKS in the circulation. Treatment with recombinant C1INH, KKS inhibitors, or B2R antagonists might be considered a long-term treatment for vascular disorders which involves neuroinflammation. Furthermore, manipulation of C1INH levels in the periphery might be used to intentionally open the BBB for drug administration.

## Materials and Methods

**Animals**. All animal experiments were conducted in accordance with the guidelines of the US NIH Guide for the Care and Use of Laboratory Animals and with approval from the Animal Care and Use Committee of The Rockefeller University. C57/Bl6J mice (males and females) were used for all experiments. A second WT model, C57/C3H, was also used. C1INH ASO (murine sequence)[18] and control ASO (CTRL ASO, no homologies to the mouse genome) were provided by Ionis Pharmaceuticals. FXII-/- mice were a gift from Thomas Renee backcrossed to C57BL/6 mice for >10 generations[83].

**ASO preparation and treatment**. C1INH ASO and CTRL ASO were dissolved in saline and injected subcutaneously to ten weeks-old mice at 150 mg/kg/week for the first two weeks (3 times a week of 50mg/kg for two weeks) and then reduced to 50 mg/kg/week for 10 weeks (twice a week of 25mg/kg for 10 weeks) (n=6-14 mice/group per cohort).

**Plasma processing**. Plasma was collected the day before the treatment started, after two weeks of treatment with 150mg/kg/week (sub-mandibular), and the day of sacrifice (12 weeks, cardiac puncture). Animals were anesthetized at the end of the treatment using regulated CO_2_. Mice were then transcardially perfused with saline. Upon collection, blood was immediately processed in EDTA-containing tubes (BD Microtainer). Heparin was avoided as it has been shown to bind to C1INH[84]. Blood was centrifuged at 1300 rpm for 15 min at room temperature (RT). The upper phase of the supernatant was transferred to a second tube containing 0.5 M EDTA PH8 and centrifuged again. The plasma was aliquoted and immediately frozen and stored at −80°C until analysis. During the lifespan of the mice, blood was taken submandibularly. At the end of the experiment, blood was taken from the heart.

**Splenocytes and peritoneal macrophages**. Spleenocytes and peritoneal macrophages were extracted from CTRL ASO- and C1INH ASO-treated mice, and single-cell suspensions were prepared to evaluate using FACS analysis [85,86]. Anti-CD4+ Cy7 conjugated and anti-CD8+ APC conjugated antibodies (BD Bioscience) were used to stain T-cells and B-cells and anti-F4/80 APC conjugated and anti-CD11b PE conjugated (BD Bioscience) were used to label macrophages. Anti-SERPING1 conjugated FITC (biorbyt orb360810) was used to detect C1INH expression in specific cells.

**Chronic restraint stress**. Chronic restraint stress (CRS) was induced [87] with slight modifications. Mice were immobilized for 6 hours using a disposable netting. Blood was collected following the restraint period, and plasma was prepared. Levels of plasma corticosterone were determined using Corticosterone Enzyme Immunoassay Kit (K014-H1, Arbor Assays).

**LPS treatment**. Mice were injected intraperitoneally (i.p.) with 100 μl of 1 mg/ml lipopolysaccharide (LPS; L-2630 Sigma-Aldrich). Blood was collected and plasma prepared 12 hours after LPS injection. This source was used as a positive control during all ELISA examinations.

**Alanine aminotransferase (ALT) activity assay**. Liver enzyme function was examined using the ALT Activity Assay (MAK052, Sigma-Aldrich). As a positive control group for this assay, we used lipopolysaccharide (LPS) to induce high levels of secreted ALT levels in the plasma[88].

**Blood pressure**. Blood pressure was measured at the end of ASO treatment using tail-cuff plethysmography (Kent Scientific). An average of 3 readings was obtained for each animal during measurement.

**Evans blue and brain edema**. Twelve hours before the sacrifice, 2% Evans blue in saline was injected into mice intraperitoneally. After perfusion with saline, one hemisphere was taken to assess percent of H_2_O volume using the wet/dry procedure [89]. Hemispheres were immediately weighed to obtain wet weight (WW) and heated to 100 °C for 24h. Then samples were weighed again to obtain the dry weight (DW). Brain water content was calculated as %H_2_O = (WW-DW) X 100/WW. The other brain hemisphere was collected and sectioned. Loss of BBB integrity was revealed by visualizing Evans blue under the fluorescence microscope.

**Immunohistochemistry**. Fresh-frozen sections were fixed with 50% MeOH and 50% acetone for 10min at −20°c. The primary antibodies used were: anti-GFAP (DAKO Z0334); anti-CD11b (Abcam ab-8878); anti-TMEM (Abcam ab209064); anti-PECAM1 (BD Pharmingen 550274); anti-CD68 (AbD Serotec MCA1957GA); anti-CD206 (Thermo PA5-46994); anti-Fibrinogen (Dako A0080); IgG (Thermo scientific); anti-laminin (Fisher scientific RT-795-PO); and anti-iNOS (Abcam ab-129372). For secondary antibodies, we used IgG from donkey anti- mouse, -rat, -rabbit or -goat, depending on the host of the first antibody (Thermo Scientific). Vectasheild-DAPI (Vector Labs) was used to seal the slides. Brain sections were visualized using Zeiss Axiovert200. Images were analyzed using ImageJ software.

**Tunnel assay**. Cell death by apoptosis was examined using the In-Situ Cell Death Detection Kit, TMR red (Roche).

**Western blot**. Western blot procedure was done as previously demonstrated [19]. Antibodies used in plasma: anti C1INH (Proteintech 12259-1-AP); anti-HK light chain (R&D MAB22061); anti-FXII (HTI PAHFFXII-s); anti-plasma kallikrein (R&D AF2498); anti-C1qA (Thermo scientific PA5-29586); anti C1r (Abcam ab66751); anti-C3a (Thermo scientific PA1-30601); anti-transferrin (Abcam ab82411). All results were normalized to transferrin levels in plasma. Brain: anti-GFAP (DAKO Z0334); anti-CD11b (Abcam ab-133357); anti-B2R (LSBio LS-C405461); anti-iNOS (Abcam ab129372); anti-oclludin (Invitrogen 33-1500); anti-GAPDH (Abcam ab9484 or Proteintech 60004-1-Ig); anti-actin (sigma A5441or proteintech 60008-1-Ig). Protein levels were quantified using NIH Image J densitometry. All results for brain protein extracts were normalized to GAPDH or Actin.

**ELISA**. ELISA was used to determine level of C3a (Molecular Innovations) and bradykinin (ENZO) in plasma and brain homogenate. The samples were normalized to 100% from CTRL-ASO to pool results from three different tests. For detection of pro-inflammatory cytokines, we used Mouse DuoSet ELISA kits for IL-1b, IL-6, and TNF-α (R&D). For a positive control in these assays, we used plasma from an LPS-injected mouse [90].

**Gene expression**. RNA was extracted from fresh cortex or frozen whole brain using RNeasy Lipid Tissue mini kit (74804, Qiagen). RNA was converted to cDNA using High Capacity cDNA Reverse Transcription Kit (4368814, Life technologies). Gene expression levels were amplified using Taqman enzyme (Applied Biosystems, 4370048) and primers (SERPING1 Mm00437835_m1, IL1-β Mm00434228_m1, IL-6 Mm00446190_m1, TNF-α Mm00443258_m1, Bdkrb1 Mm04207315_s1, Bdkrb2 Mm00437788_s1, C3ar1 Mm01184110_m1, PECAM1 Mm01242576_m1, GFAP Mm01253033_m1, CD11b Mm00434455_m1, and normalized to either endogenous mouse GAPDH Mm99999915_g1 or ACTB Mm02619580_g1. ΔΔCT was quantified and compared between samples.

**Kallikrein activity assay**. Plasma of only males was evaluated for KKS activity using the chromogenic substrate S-2303 (Diapharma) or Pefachrome PK8092 (Pentapharm) as previously demonstrated[19].

**Behavioral analysis**. All behavioral experiments were performed and analyzed with a researcher blinded to genotype and treatment. Fear conditioning was performed as previously described [91] with slight changes. Two foot shocks were given (0.7 milliamp, 0.5 sec), one after the first three minutes and the second at the end of the five minutes of the first day. Open field test was done as previously described [92].

**Statistical analysis**. All statistical analyses as were determined by two-tailed t-test. One-tailed t-test was performed when the results confirmed the hypothesis. Some results were compared to total 100% of CTRL ASO (dotted line) and numerical values presented in graphs are mean ±SEM. Results that were mean ± 2*SD were excluded.

## Acknowledgements

The authors thank members of the Strickland laboratory for their help. This work was supported by The EGL foundation and NIH; Rudin Family Foundation; Mellam Family Foundation; Mr. John A. Herrmann Jr.; and Mary & James G. Wallach Foundation.

## Authorship

Contributions: D.F. designed the study, performed experiments, analyzed data, and wrote the manuscript; E.F. and A.R. performed experiments and provided technical assistance; E.H.N and S.S participated in data analysis and manuscript preparation. Conflict-of-interest disclosure: Alexey S. Revenko and A. Robert MacLeod are employees and stockholders of Ionis Pharmaceuticals. The authors declare no additional competing financial interests.

---

**Supp. Fig. 1.**
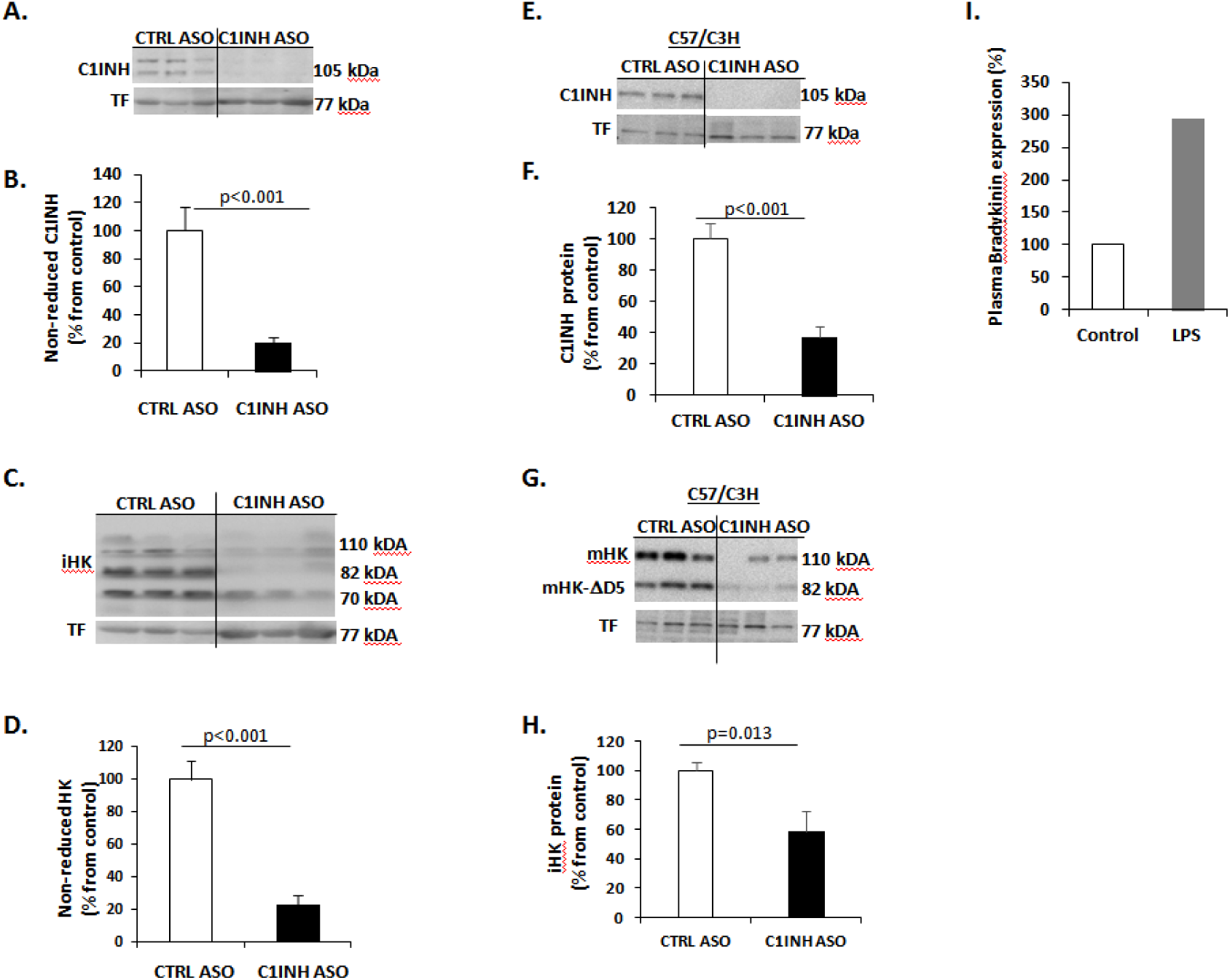
Depleted circulating C1INH in non-reduced conditions and in C3H/C57 C1INH ASO treated mice. **(A)** Representative western blot of non-reduced plasma probed with anti-C1INH. **(B)** Quantification of C1INH expression levels between CTRL ASO and C1INH ASO (p<0.001, n=6) **(C)** Representative western blot of non-reduced plasma probed with anti-iHK. **D)** Quantification of iHK expression levels between CTRL ASO and C1INH ASO (p<0.001, n=6). **(E)** Representative western blot of reduced plasma from C57/C3H mouse model treated with C1INH ASO probed with anti-C1INH. **(F)** Quantification of plasma from C57/C3H mouse model treated with C1INH ASO showed a decreased iHK expression compared to CTRL ASO (p<0.001, n=6). **(G)** Representative western blot of reduced plasma from C57/C3H mouse model treated with C1INH ASO probed with anti-iHK. **(H)** Quantification of plasma from C57/C3H mouse model treated with C1INH ASO showed a decreased iHK expression compared to CTRL ASO (p=0.013, n=6). **(I)** Bradykinin levels in LPS treated mice measured by ELISA showed 3-fold increase.

**Supp. Fig. 2.**
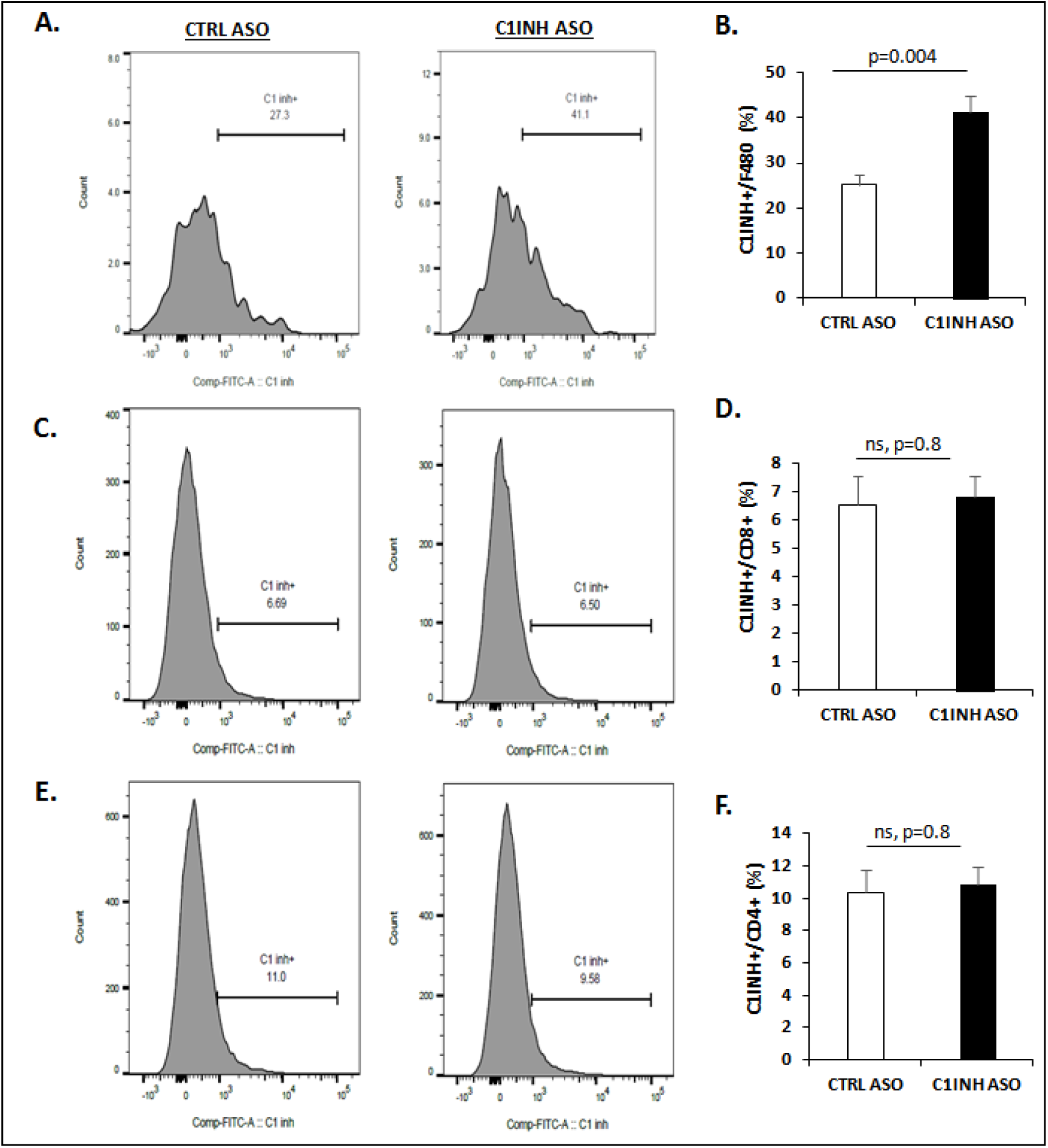
C1INH expression levels are not reduced by the ASO treatment in other immune cell types. **(A)** Representative histograms between CTRL ASO and C1INH ASO showing C1INH expression gated from positive cells expressing F4/80. **(B)** Quantification of the histograms showing increased expression (p=0.004, n=5/group) of C1INH in F4/80 macrophages. **(C)** Representative histograms between CTRL ASO and C1INH ASO showing C1INH expression gated from positive cells expressing CD8+ cells. **(D)** Quantification of the histograms showing No differences in C1INH expression (p=0.8, n=5/group) in CD8+ splenocytes. **(E)** Representative histograms between CTRL ASO and C1INH ASO showing C1INH expression gated from positive cells expressing CD4+ cells. **(F)** Quantification of the histograms showing No differences in C1INH expression (p=0.8, n=5/group) in CD4+ splenocytes.

**Supp. Fig. 3.**
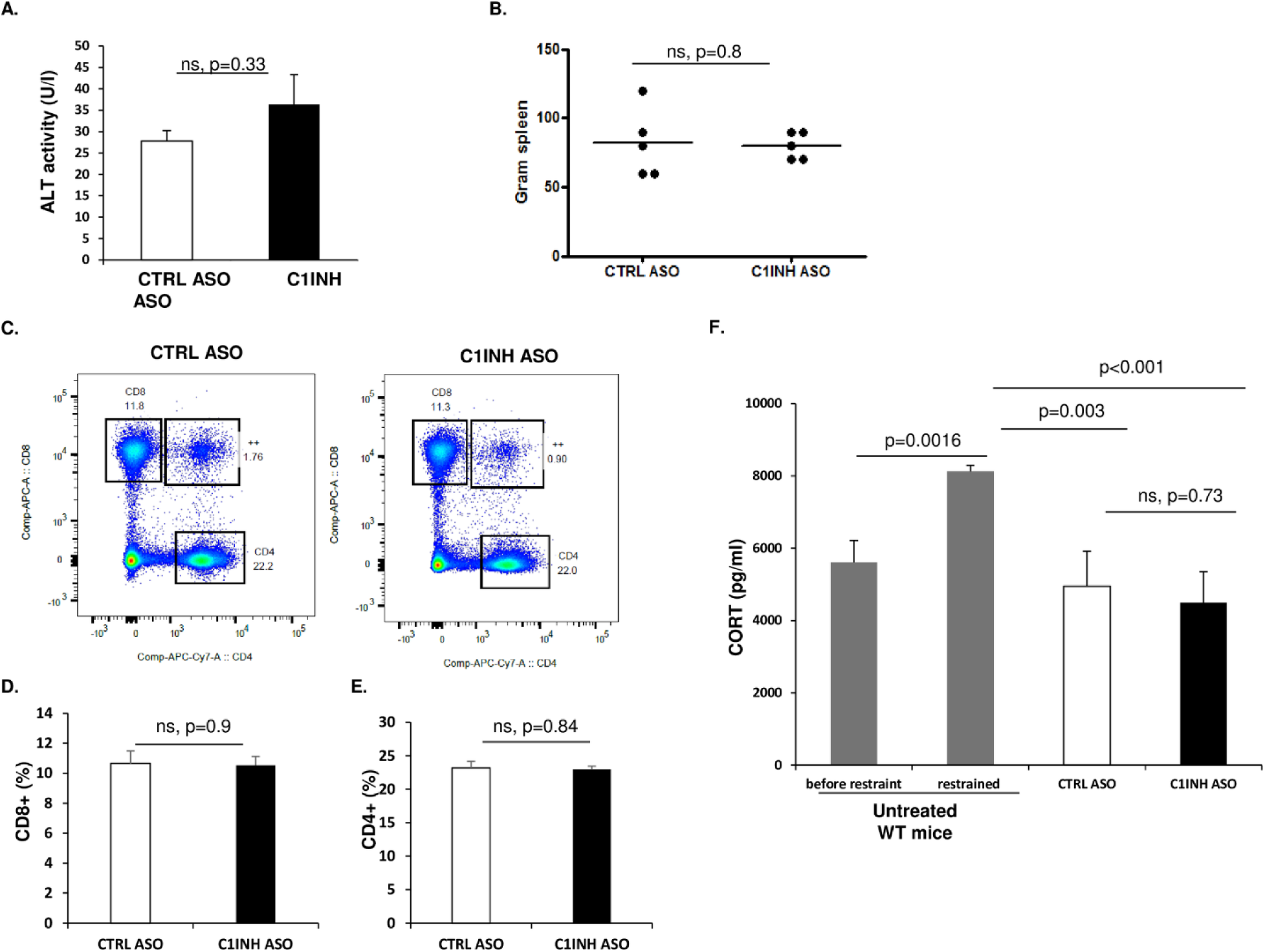
Knock-down of circulating C1INH did not induce toxicity. **(A)** Activity level of liver function measured by alanine transferase (ALT) assay in mouse plasma of CTRL ASO treated mice vs. C1INH ASO mice (27.8±2.41 vs. 36.34±6.98, p=0.33; n=4–5/group). **(B)** Spleen weight measurement after treatment showed no difference between CTRAL ASO and C1INH ASO (82±11.14 vs. 80±4.47, p=0.87, n=5). **(C)** Representative FACS dot-plot of positive CD4+ and CD8+ from splenocytes of CTRL ASO vs. C1INH ASO treated groups. **(D and E)** Quantification analysis of CD8+ and CD4+ cells show no differences in activation levels (10.66±0.83 vs. 10.53±0.59, p=0.9, and 23.18±1.01 vs. 22.96±0.48, p=0.84, n=5). **(F)** Corticosterone levels did not differ (p=0.73, n=5) between treated groups. For positive control we used 6 hours-restrained mice compared to levels of CORT before restraining (p=0.001, n=7).

**Supp. Fig. 4.**
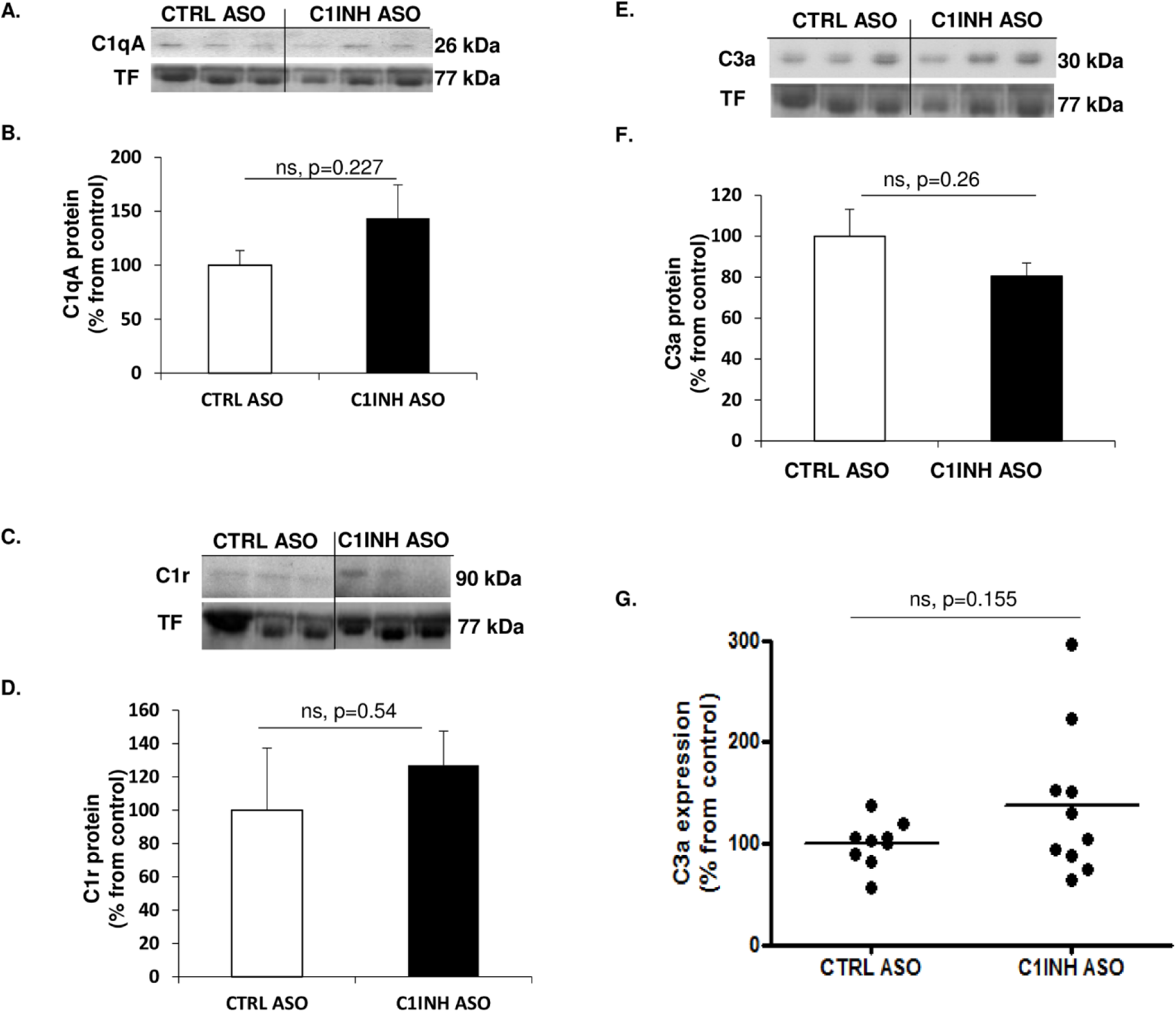
Depletion of C1INH did not induce complement activation. **(A)** Representative western blot of reduced plasma probed with anti-C1qA. **(B)** Quantification of C1qA expression levels show no difference between CTRL ASO and C1INH ASO (p=0.227, n=6). **(C)** Representative western blot of reduced plasma probed with anti-C1r. **(D)** Quantification of C1r expression levels show no difference between CTRL ASO and C1INH ASO (p=0.54, n=6). **(E)** Representative western blot of reduced plasma probed with anti-C3a. **(F)** Quantification of C1r expression levels show no difference between CTRL ASO and C1INH ASO (p=0.26, n=6). **(G)** ELISA of C3a levels show no difference in plasma of CTRL ASO vs. C1INH ASO (p=0.155, n=9–10/group).

**Supp. Fig. 5.**
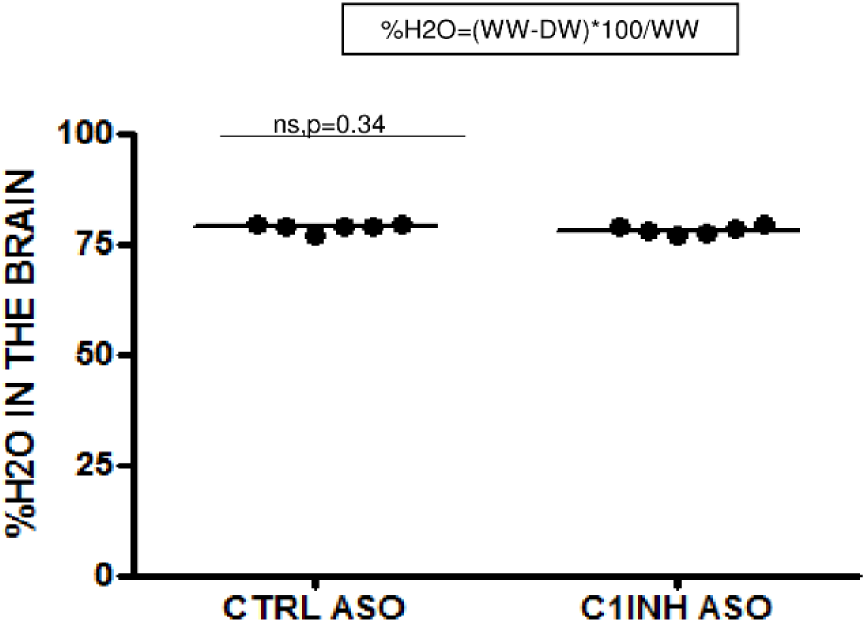
**Measurement of water content in the brains of treated mice showed no differences** (p=0.34, n=6).

**Supp. Fig. 6.**
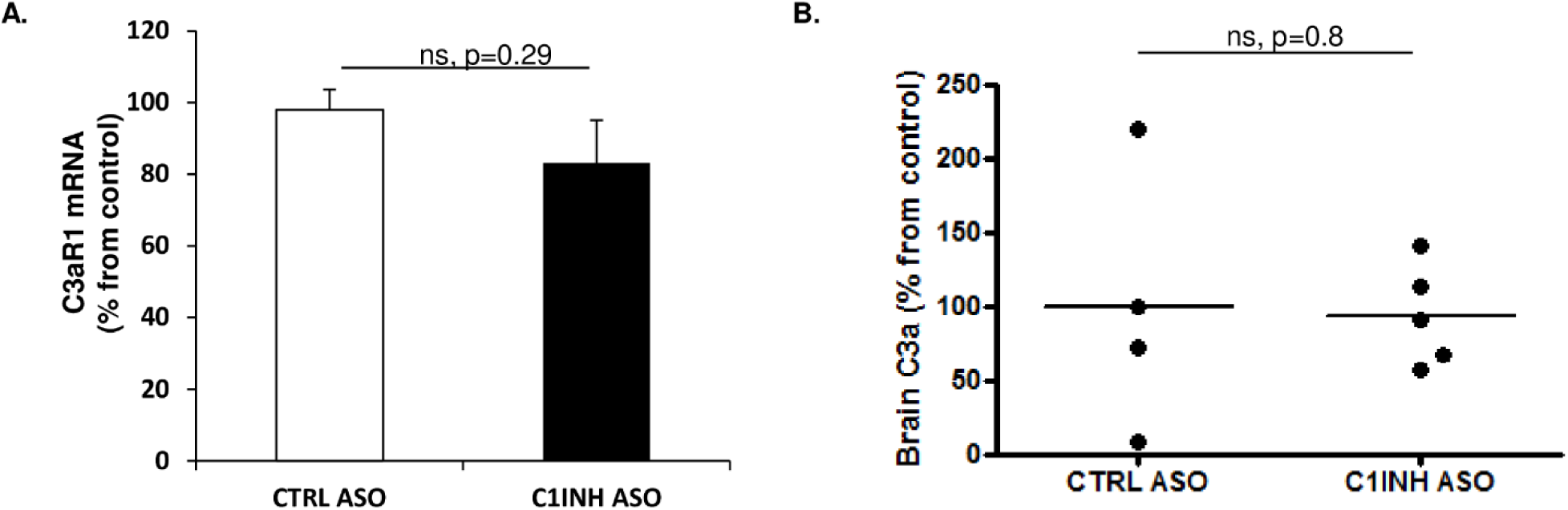
No changes in gene or protein expression of complement system C3a and C3aR1. **(A)** No changes were found in the gene expression levels of C3aR1 in the brains of C1INH ASO treated mice compared to control mice (p=0.29, n=6/group). **(B)** No differences were detected in C3a protein expression in brains of C1INH ASO vs. CTRL ASO mice (p=0.8, n=4–5).

**Supp. Fig. 7.**
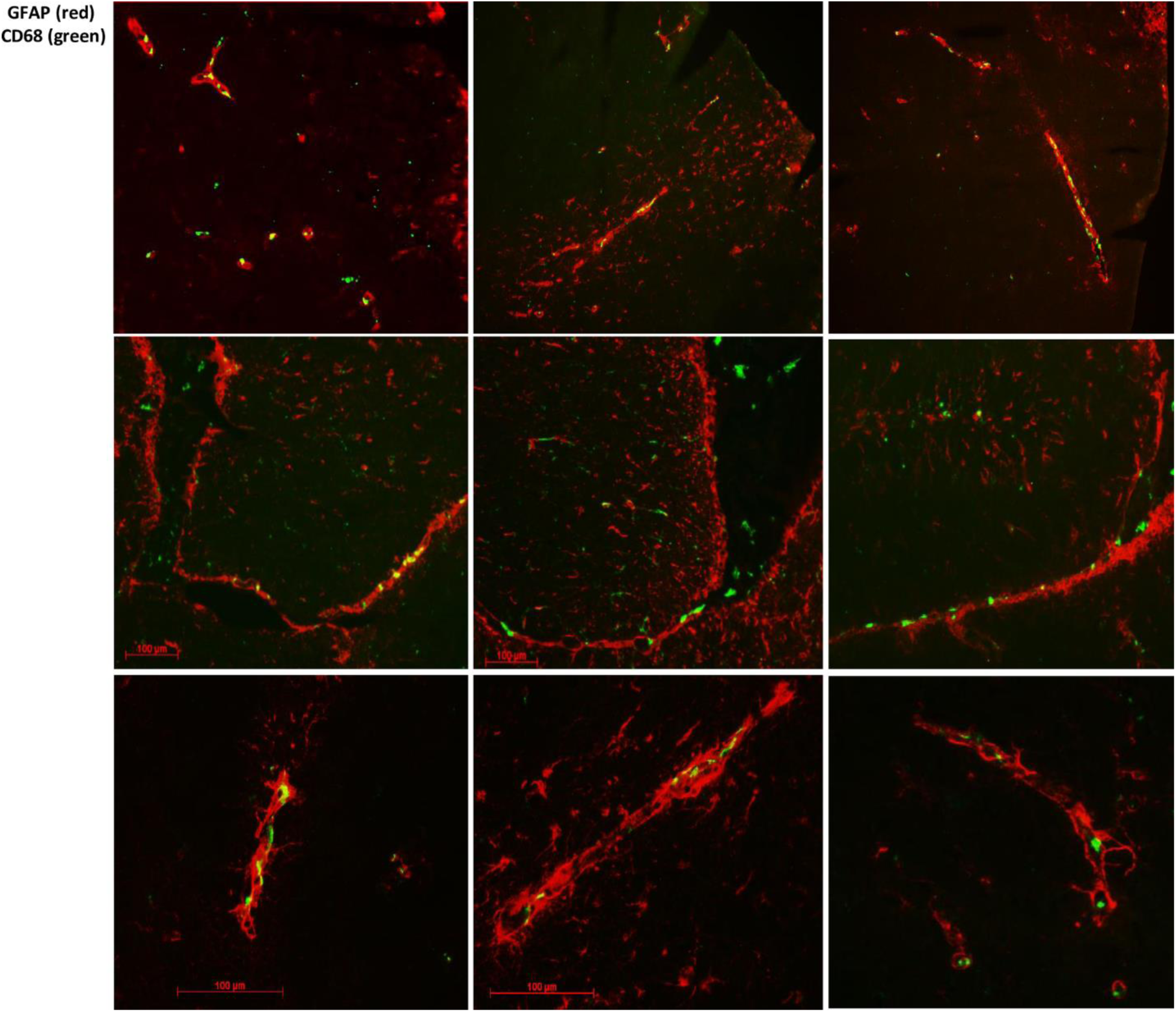
Representative images of GFAP (red) staining and CD68 (green) in the neurovascular units.

**Supp. Fig. 8.**
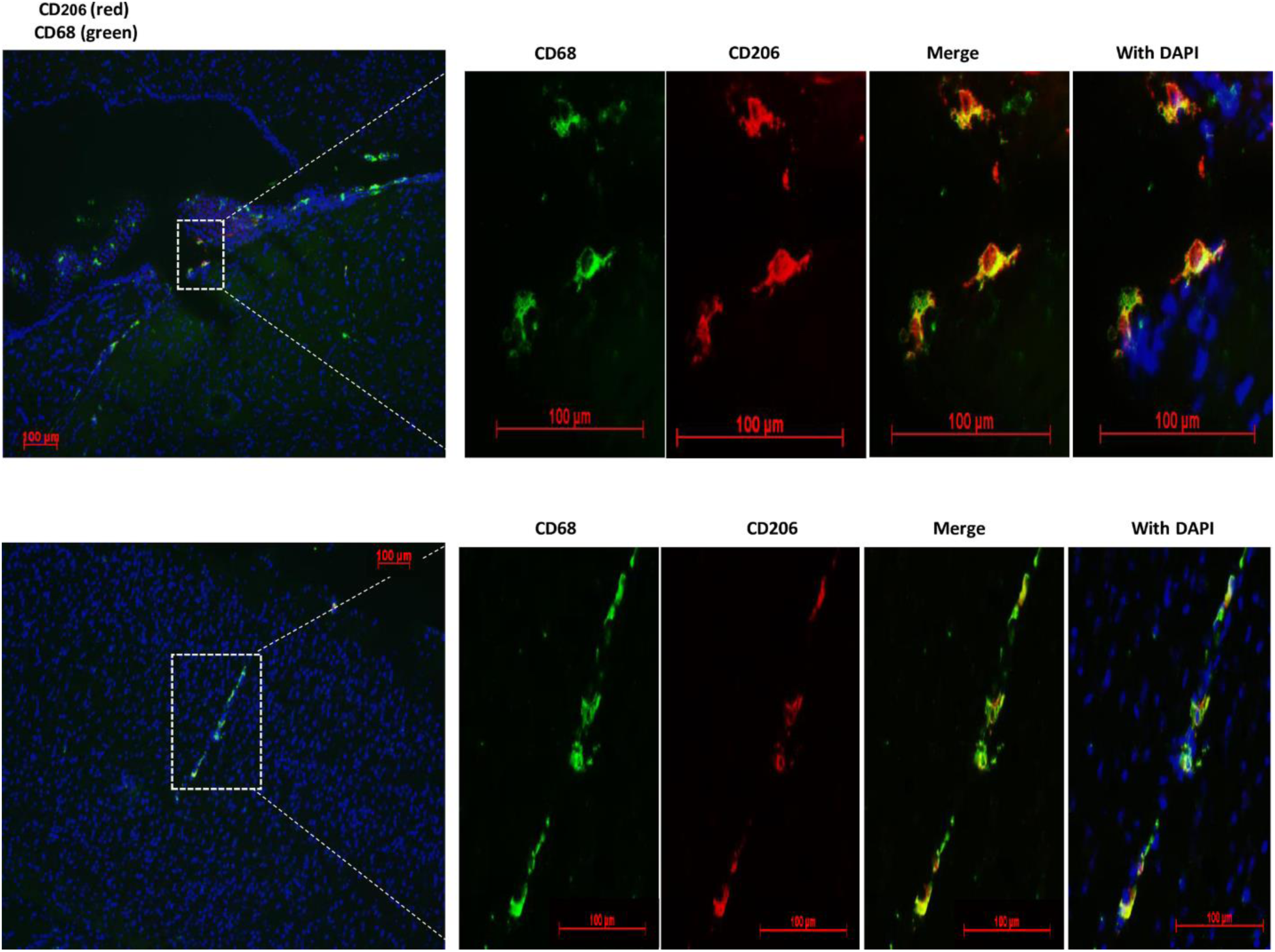
Co-localization of CD68 positive cells and CD206.

**Supp. Fig. 9.**
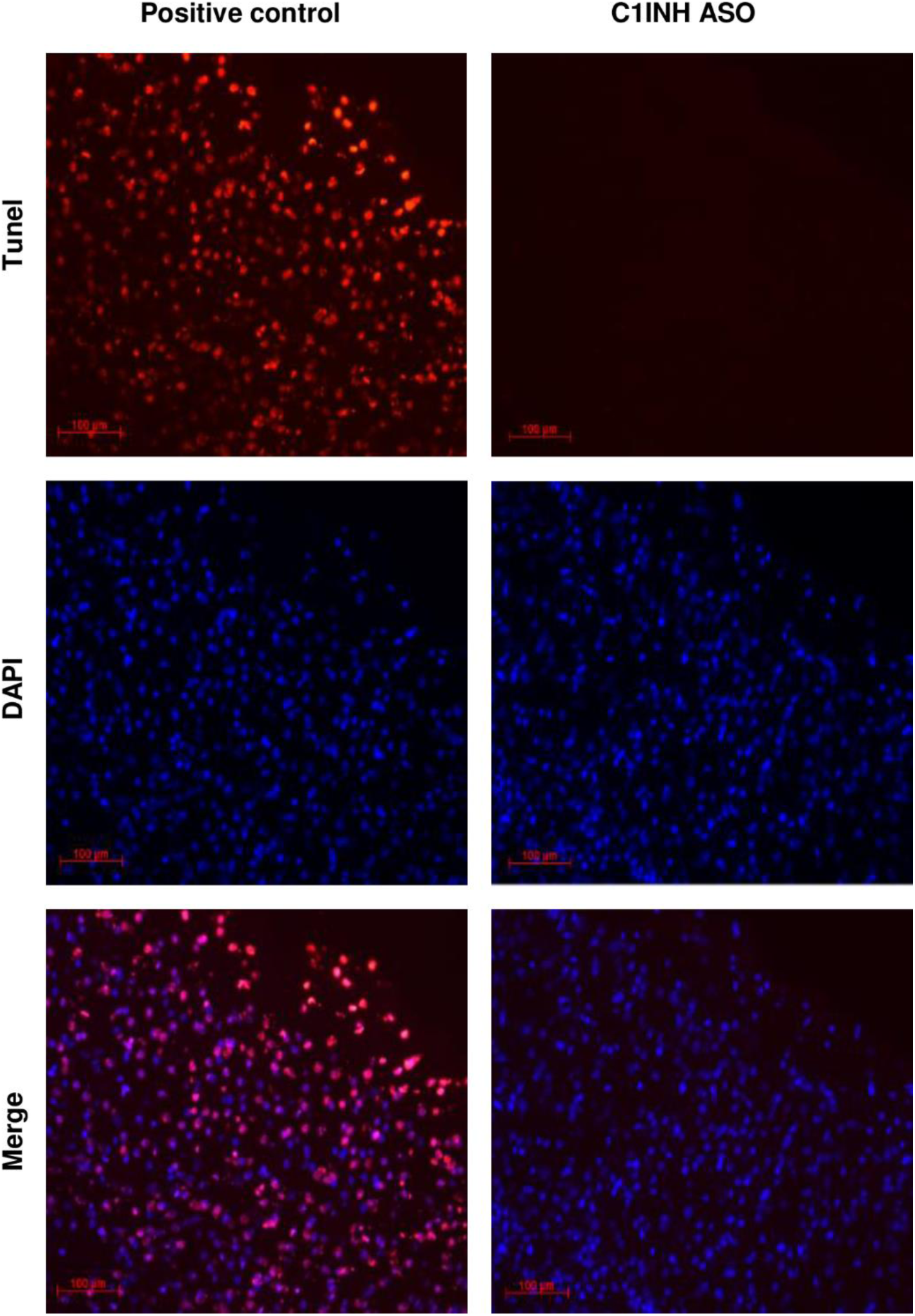
Negative Tunel staining in brains of C1INH ASO treated mice. Representative image of Tunel staining showing no positive Tunel staining in the brains of C1INH ASO treated mice indicating non-apoptotic cells induced.

